# A Pilot Study of Transcranial Magnetic Stimulation and the Right *Teaching* Parietal Junction

**DOI:** 10.1101/2025.09.15.676361

**Authors:** Molly Skowron, Ashton Morante, Alexa Alvarez, Julian Paul Keenan

## Abstract

Theory of Mind (ToM) is conceptualized as the ability to infer the thoughts and feelings of others. Many studies have found that ToM abilities play a central role in collaborative communication between adults, with teachers performing more effectively when they have high levels of ToM. One of the main brain regions implicated in ToM is the right temporoparietal junction (rTPJ), which is thought to be responsible for evaluating a person’s belief formation process. The current study had participants engage in a Lego model building task consisting of a director who instructed a builder on how to create duplicate models from a prototype that only the director could see. The rTPJ of the director was targeted for excitatory (10 Hz) transcranial magnetic stimulation (TMS) and inhibitory (1 Hz) TMS. There was a trend such that there was a decrease in the amount of time needed and an increase in accuracy between excitatory TMS and sham TMS (the control). The opposite pattern was observed between inhibitory TMS and sham TMS. These results suggest that teaching abilities are likely at least partially dependent on the rTPJ.

---

Common ground is defined as the knowledge or beliefs that are shared between the two individuals who are attempting to communicate (Clark et al., 1983). In order for individuals to exchange information cooperatively, they must establish a common ground to adjust both their verbal and non-verbal communication (Clark & Brennan, 1991). Theory of Mind (ToM) is conceptualized as the ability to infer the thoughts and mental states of others (Keenan et al., 2003; Premack & Woodruff, 1978).

It has been suggested that a high level of Theory of Mind (ToM) may be associated with an individual’s ability to establish common ground, leading to fewer misunderstandings during collaborative communication (Gupta et al., 2012; Rubio-Fernandez, 2021). It is possible that people do not always fully use their ToM abilities in tasks that are mentally demanding such as speaking (Keysar et al., 2003). Keysar concluded that an adult’s ability to represent others’ beliefs does not reliably interpret their behavior. Furthermore, while adults are able to efficiently switch perspectives, they fail in effectively using other people’s knowledge to interpret what they mean (Apperly et al., 2010). However, some skeptics believe these findings are inaccurate measurements of ToM since some evidence suggests that the methods used in these studies are actually measuring a participant’s selective attention, rather than their ToM abilities (Rubio-Fernandez, 2017).

Meanwhile, many studies have found that ToM abilities play a central role in collaborative communication between adults. Champagne-Lavau and colleagues confirmed that individuals adapt their communication styles based on who they are communicating with, whereas those with ToM deficits individuals (such as autism and schizophrenia) were not able to in the same way (2009). A different study demonstrated that individuals’ ToM abilities were linked to their ability to adjust their communication style to facilitate understanding with the individual they were communicating with (Achim et al., 2015).

## 1.1. The Role of Theory of Mind in Teaching

Previously, we conducted a study in which pairs of participants engaged in a Lego model building task consisting of a director who instructed a builder on how to create duplicate models from a prototype that only the director could see instructions to (Krych-Appelbaum et al., 2007). This study demonstrated that a high level of ToM as demonstrated by the Mind in the Eyes (MIE) test was an advantage when instructing, resulting in fewer building errors, but was a *disadvantage* when following. The MIE is a more advanced test of ToM that has participants looking at photographs of people’s eyes, and best trying to describe the emotion they are feeling based on two words (Baron-Cohen & Jolliffe, 1997). In other words, people in the role of director with a high level of ToM may be better at noticing when their partner is confused and may be more likely to adjust their instruction accordingly to minimize misunderstandings (Achim et al., 2015).

However, in the role of the builder, a high level of ToM was a disadvantage, as it is possible that builders with higher ToM may *assume* they know what their partner means instead of asking for clarification, resulting in errors in communication. It has been shown that comprehension in an interview is most accurate when an interviewer can anticipate when their respondent has not understood a question, and when a respondent asks questions to clarify their understanding (Schober et al., 2004; Suessbrick et al., 2000). It is possible that high levels of ToM are an advantage for a teacher due to the nature of classroom instruction. While teaching, the teacher is expected to create a mental model of how the student might respond to various teaching choices and how their learning might be affected by their emotions (Rodriguez, 2013). Teachers must then use both a metacognitive understanding of the behavior or task they are teaching and their mental model of the student’s mind (Csibra & Gergely, 2006).

## 1.2. Theory of Mind and the Right Temporoparietal Junction Teaching

Numerous functional magnetic resonance imaging (fMRI) studies have implicated the right temporoparietal junction (rTPJ) in ToM tasks such as false belief tasks (Boccadoro et al., 2019; Dodell-Feder et al., 2011; Mossad et al., 2016), anticipating an opponent’s move in chess (Powell et al., 2017), and attribution of intentions to others (Völlm et al., 2006). The rTPJ was also implicated in ToM tasks in studies which used functional near-infrared spectroscopy (fNIRS) (Hyde et al., 2015). Two developmental fMRI studies confirmed that the functional specialization of the rTPJ was associated with improved performance on ToM tasks (Gweon et al., 2012; Mukerji et al., 2019). However, the rTPJ does not complete ToM tasks on its own. Other brain regions have also been shown to be associated with ToM tasks, such as the medial prefrontal cortex (MPFC), superior temporal sulcus (STS), and precuneus (PC) (Carrington & Bailey, 2009; Saxe & Powell, 2006). Each of these regions is thought to play a distinct role in representing the mental states of others, and evidence suggests the rTPJ is responsible for containing information about what another person knows or should know and for evaluating a person’s belief formation process (Koster-Hale et al., 2017).

While some authors have suggested that rTPJ is only involved in the preliminary social cognition processes that aid in ToM rather than being responsible for ToM reasoning itself (Gallagher & Frith, 2003), other authors have directly refuted this by demonstrating that the rTPJ was active when participants were reasoning about the mental states of another person, and the presence of that person alone without the reasoning was not enough to result in rTPJ activation (Saxe & Kanwisher, 2013). Another hypothesis that has been suggested is that the rTPJ plays a role in reorienting a person’s attention to an unexpected stimulus, and its activation during ToM tasks is coincidental due to the presence of an unexpected stimulus (Buccino et al., 2007).

Several authors have provided evidence to refute this hypothesis, finding that there are two nearby but distinct regions, one of which is activated by ToM tasks and the other is involved in the reorienting of attention to unexpected stimuli (Krall et al., 2015; Scholz et al., 2009). Further research confirmed that there was a significant difference in rTPJ activation for tasks involving ToM when compared to non-ToM tasks, and this difference was not found when comparing expected and unexpected stimuli (Young, Dodell-Feder, & Saxe, 2010). However, there is still controversy about these findings, as other studies have found evidence suggesting that there is a shared cognitive mechanism for attention and ToM in the rTPJ, rather than a separation of the two (Schuwerk et al., 2021).

Several studies have demonstrated that Transcranial Magnetic Stimulation (TMS) applied to the rTPJ directly influences an individual’s ToM abilities. Bardi et al. (2017) applied TMS to inhibit the rTPJ while the individual was engaged in a spontaneous ToM task. Their evidence suggested that the participants’ ability to represent the beliefs of others was unaffected, but their ability to predict future events based on this knowledge was strongly impacted. Other studies have found that inhibitory brain stimulation to the rTPJ caused participants to take the beliefs of others into consideration at a much lower rate, specifically in regard to the mental state reasoning necessary to make moral judgements on the actions of others (Chou & Chen, 2021; Young, Camprodon, et al., 2010). The opposite effect was observed when transcranial direct current stimulation (tDCS) was used to increase activity in the rTPJ. While participants with an inhibited rTPJ did not take an individual’s mental state into account (they were more likely to see no moral issue with an individual who intended to harm another but failed), participants with increased rTPJ activity were more likely to take an individual’s mental state into account (they were more likely to see no moral issue with an individual who harmed another by accident, with no ill intent) (Sellaro et al., 2015). A study by Filmer et al. (2019) found that tDCS applied to the rTPJ affects an individual’s ability to engage in traditional ToM tasks such as the false belief task.

Two psychiatric disorders which are frequently associated with ToM deficits are schizophrenia and autism. Several neuroimaging studies have implicated the rTPJ as a region which is associated with the ToM deficits observed in these individuals (Bitsch et al., 2019; Brunet-Gouet & Decety, 2006; Vucurovic et al., 2020). Brain stimulation to the rTPJ has been shown in some instances to affect symptoms of schizophrenia (Lee et al., 2005), but these results have not been consistently replicated with specific regard to ToM symptoms (Klein et al., 2021).

Autism has also been associated with ToM deficits (Baron-Cohen et al., 1985). Several neuroimaging studies have linked rTPJ functioning to ToM deficits in individuals with autism (Dichter, 2022; Kana et al., 2015; Nijhof et al., 2018). Brain stimulation to the rTPJ has also been shown to consistently affect ToM in individuals with autism (Esse Wilson et al., 2018; Salehinejad et al., 2021), which suggests a causal relationship between rTPJ functioning and ToM abilities in individuals with autism. One study suggested that there were no long term effect in improved ToM in participants with autism after undergoing ToM interventions that aimed to teach participants the component skills of ToM (Fletcher-Watson et al., 2014).

Since ToM plays such a big role in teaching (Achim et al., 2015; Champagne-Lavau et al., 2009; Krych-Appelbaum et al., 2007), it logically follows that brain regions responsible for ToM such as the rTPJ would be implicated in teaching as well (Filmer et al., 2019; Koster-Hale et al., 2017). Another brain region which has been associated with perspective-taking and ToM tasks is the MPFC (Carrington & Bailey, 2009; Hillebrandt et al., 2013; Saxe & Powell, 2006; Van Overwalle & Baetens, 2009). Therefore, it follows that a teacher likely uses both the rTPJ and the PFC while teaching. Vélez et al. (2023) supported this by demonstrating that when teachers are choosing which examples to use in a lesson, they choose examples that will maximize a learner’s belief in a target concept, while also taking their own preferences and communicative costs into account. The brain regions that were implicated by this study were the bilateral TPJ and the dorsal and middle MPFC (Vélez et al., 2023). A study by Zheng et al. (2018) discovered that the teacher’s rTPJ is involved in interpersonal neural synchronization (INS) during teacher-student interactions, and that the region that it synchronizes with in the brain of the student is the anterior superior temporal cortex (aSTC). Previous studies have implicated the aSTC in the representation of semantic knowledge (Correia et al., 2014; Pobric et al., 2016). This study found that when brain activity of the teacher at the rTPJ preceded that of the student at the left aSTC, a significant increase in INS was positively correlated with student levels of learning, which the researchers proposed reflected the teacher’s rTPJ predicting the mental state of the student (Zheng et al., 2018).

In the current study, we suspect that ToM abilities will be influenced by TMS applied to the rTPJ, which will in turn affect their ability to engage in the Lego model building task. When TMS is used to excite the rTPJ of an individual, they should be able to instruct faster and with increased accuracy. Therefore, the trial with the lowest average build time per model and the highest accuracy should be when the director’s rTPJ has been excited, and the reverse should occur when the director’s rTPJ has been inhibited.

## 2. Methods

### 2.1. Participants

A total of 5 participants were recruited as per local IRB guidelines via social media, email, flyers, and by word of mouth. The individuals that were recruited passed an initial safety screening questionnaire, and were only required to attend one session, lasting approximately one hour. All participants provided informed consent and were treated in accordance with guidelines set forth by t e Internal Review Board at Montclair State University and guidelines of the American Psychological Association. All TMS was delivered within the parameters provided by Wassermann (1998). As per local IRB guidelines, all TMS was delivered by the PI (approved by Montclair State University IRB).

### 2.2. Experimental Setup and Design

The director and builder sat on opposite sides of a table. There was a low barrier on the table which prevented the builder from seeing the director’s prototype but allowed the director to see the builder’s workspace. The barrier did not prevent the pair from seeing one another’s faces. The builder had access to a box containing dozens of assorted blocks (Figure 1).

**Figure 1.**
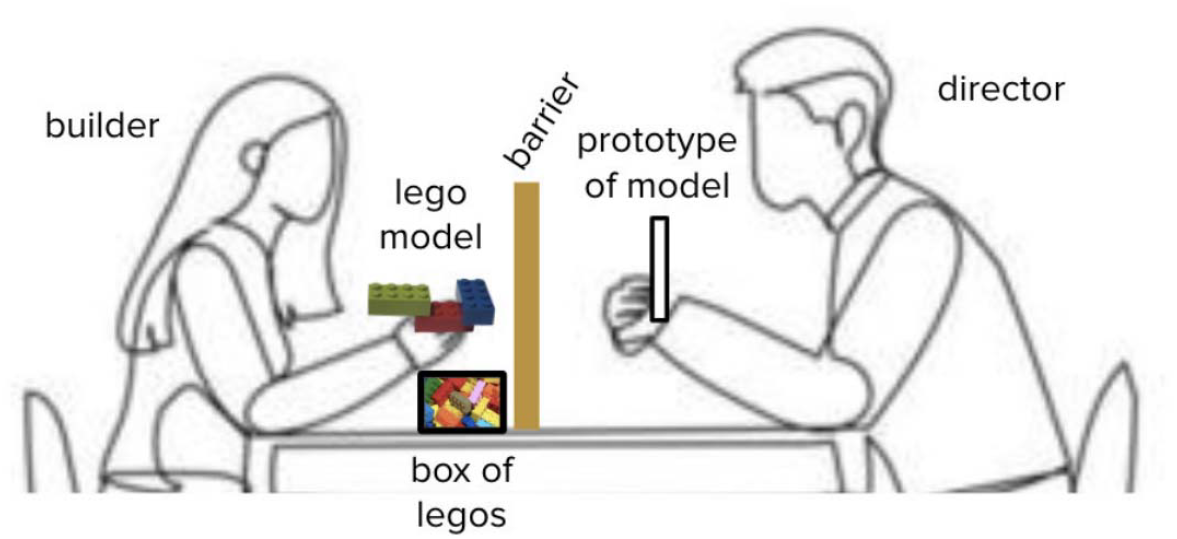
Illustration of experimental setup.

The 26 Lego prototypes each consisted of three to four 2×4 Lego blocks two to three blocks high. They were designed so they could not be simply described as familiar objects such as bridges, animals, or buildings. Pilot testing showed that the models took roughly equal time to assemble. The models provided for each experimental condition were randomized, and the participants never built the same model twice.

### 2.3. Procedure

The participant acted as the director whose role was to describe for the builder how to create duplicate models of Lego blocks, and a research assistant acted as the builder. The director was able to talk as much as needed and gesture, while the builder was told not to ask questions.

The pair was given four practice models to orient them to the task with no TMS applied. They were told to create an exact replica of each of the director’s prototypes. Each prototype was only visible to the director. After each model was completed, the pair would indicate that they were finished, then the experimenter would give them feedback by displaying the participants’ model and the original prototype before proceeding to the next model.

Afterwards, the participant was taken to receive their first round of TMS. A total of three TMS trials were conducted: excitatory TMS to the director’s rTPJ, inhibitory TMS to the director’s rTPJ, and sham TMS to the director’s rTPJ. The order of these three conditions was randomized for each participant. Following TMS application, the participant was brought back to the Lego building room. The experimenter gave them six prototype images, and instructed the pair that they would have four minutes to construct as many of the models as possible. Just like the practice round, the director with the prototype images was to give instructions to the builder who could not see the prototype images, and they would try to have the builder construct an exact replica of each prototype in the picture. The experimenter then began a four-minute timer and instructed the pair to begin. When they finished a model, the experimenter would pause the timer and give time for feedback before deconstructing the model and restarting the timer when the participant indicated they were ready. When the timer was completed or when all the models were complete (whichever occurred first), the experimenter instructed the pair to stop. This procedure was followed for each trial. The accuracy of each Lego model and the amount of time needed to complete each model was measured as an approximation of how well the director can give instructions.

### 2.4. Transcranial Magnetic Stimulation Procedure

For all TMS, a Magstim 200 Rapid pulse 1.5T and a 7 cm figure-of-eight coil were used to deliver pulses at 10 hertz (Hz) and 1 Hz. Motor Threshold via Motor Evoked Potentials (MEPs) was first established for each participant using Trigno wireless MEP amplifiers running DelSys software. The MEP was the minimal amount of stimulation output needed to induce a motor evoked response in 5 out of 10 trials (Chail et al., 2018). Participants received 300 pulses of 1Hz TMS stimulation to inhibit the rTPJ, or they received 300 pulses of 10Hz TMS stimulation to excite the rTPJ (Figure 2). The participants wore both earplugs and Lycra swim caps for the duration of the experiment.

**Figure 2.**
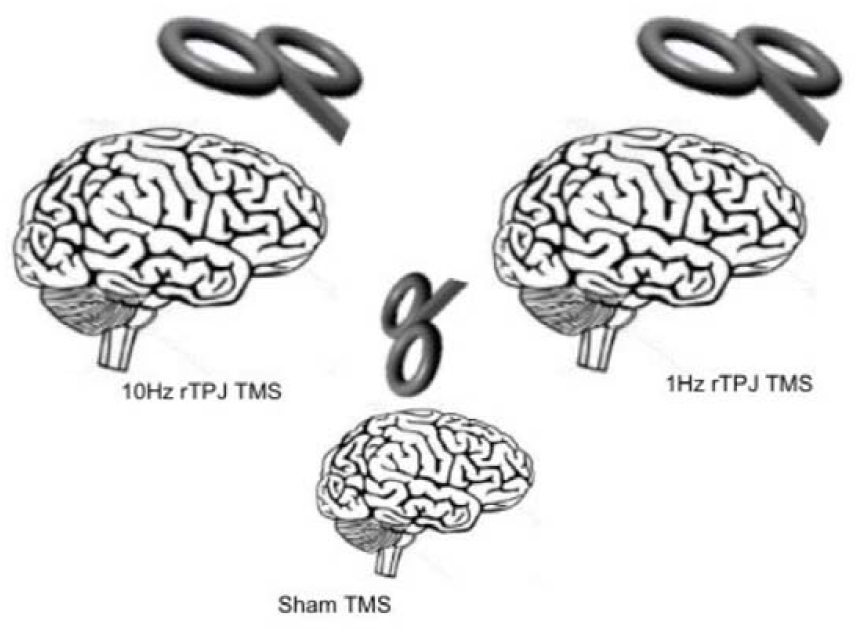
Overall TMS procedure. Participants received TMS 3 times, in a random order.

### 2.5. Statistical Analysis

We performed a one-way repeated measures ANOVA to determine if there was a significant difference (all comparisons at 0.05) in the average time to complete a model and the average model accuracy for each TMS condition.

## 3. Results

We first examined the average time taken to complete a Lego model and tested the hypothesis that excitatory TMS decreases time taken and inhibitory TMS increases time taken by using a one-way repeated measures ANOVA. It was found that there is no significant difference (F = 1.03, p = 0.39; df = 2; Figure 3). Despite the lack of statistical significance, we did notice a trend between brain stimulation and time needed to complete a model in which excitatory TMS reduced the amount of time needed and inhibitory TMS increased the amount of time needed when compared to the baseline sham condition.

**Figure 3.**
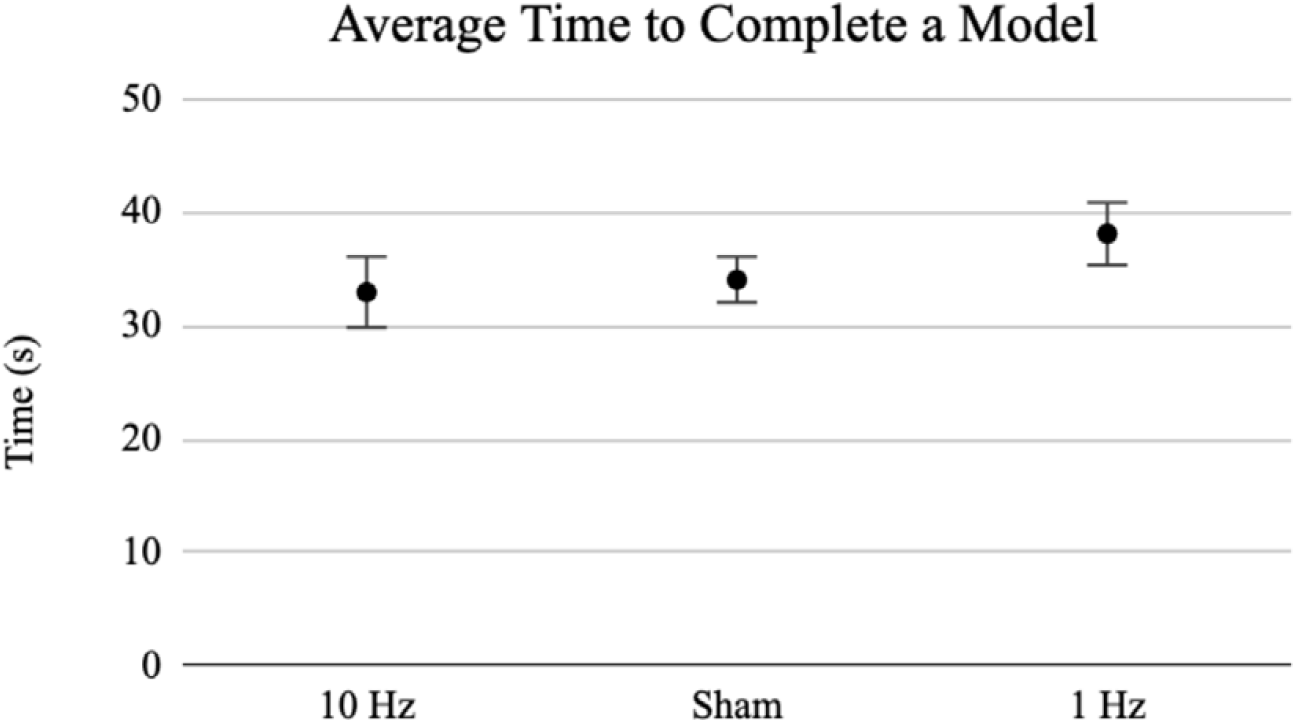
Average time to complete a Lego model for each TMS condition. All TMS conditions were found to be nonsignificant (p > 0.05). Standard errors of the means are plotted.

We then examined the average model accuracy using a one-way repeated measures ANOVA and found that brain stimulation did not result in a significant difference (F = 0.84, p = 0.45; df = 2; Figure 4). There was a trend between brain stimulation and model accuracy in which excitatory TMS increased the accuracy of the models when compared to the baseline sham condition.

**Figure 4.**
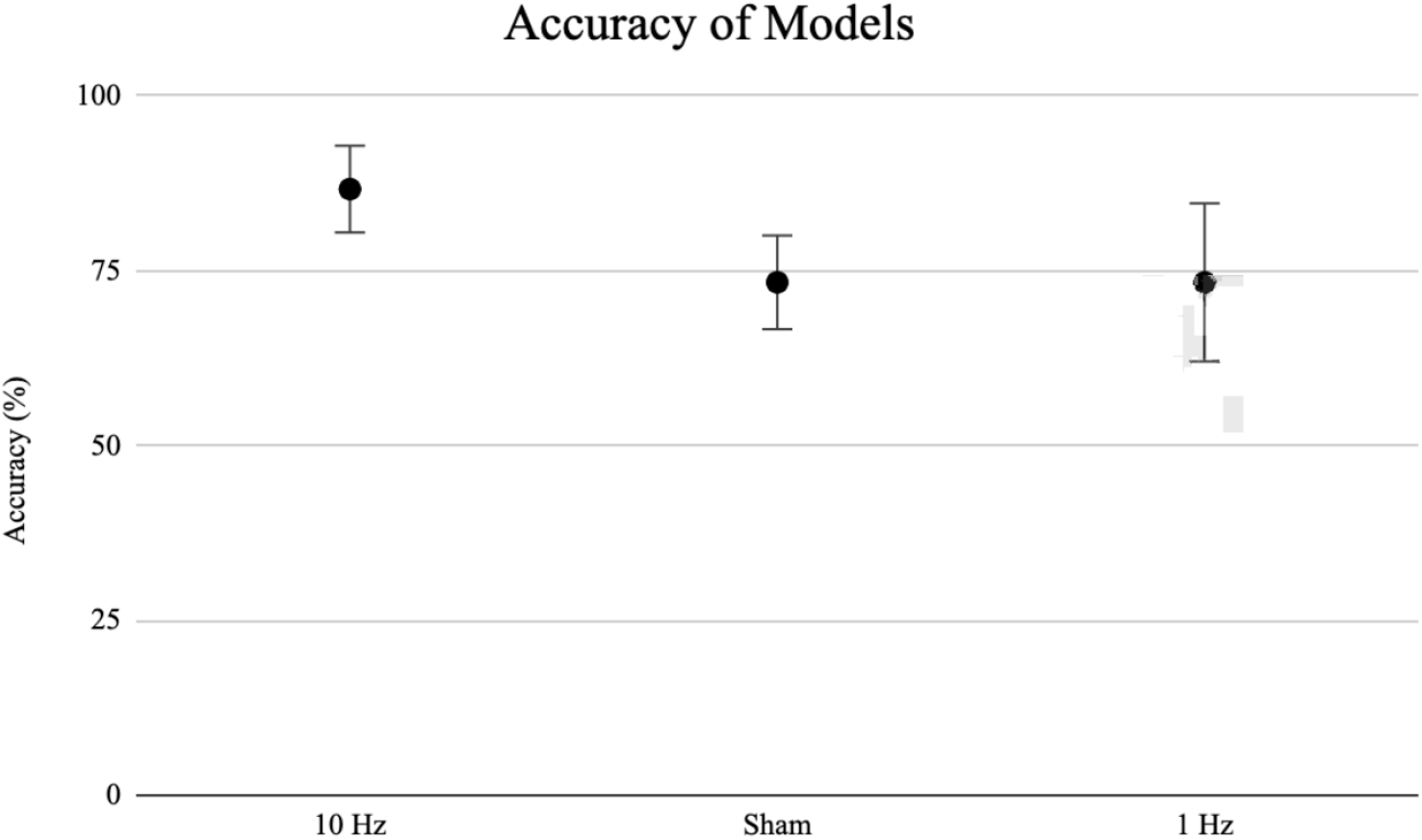
Average percent accuracy of the Lego models for each TMS condition. All TMS conditions were found to be nonsignificant (p > 0.05). Standard errors of the means are plotted.

We then multiplied time in seconds by the number of mistakes made to create a combined measure of speed x accuracy for each trial. We compared these values using a one-way repeated measures ANOVA and found that brain stimulation did not result in a significant difference (F = 0.84, p = 0.46; df = 2; Figure 5). Despite the lack of statistical significance, a trend emerged between brain stimulation and combined speed/accuracy in which excitatory TMS resulted in improved speed/accuracy of the models when compared to the baseline sham condition, and inhibitory TMS resulted in worse performance.

**Figure 5.**
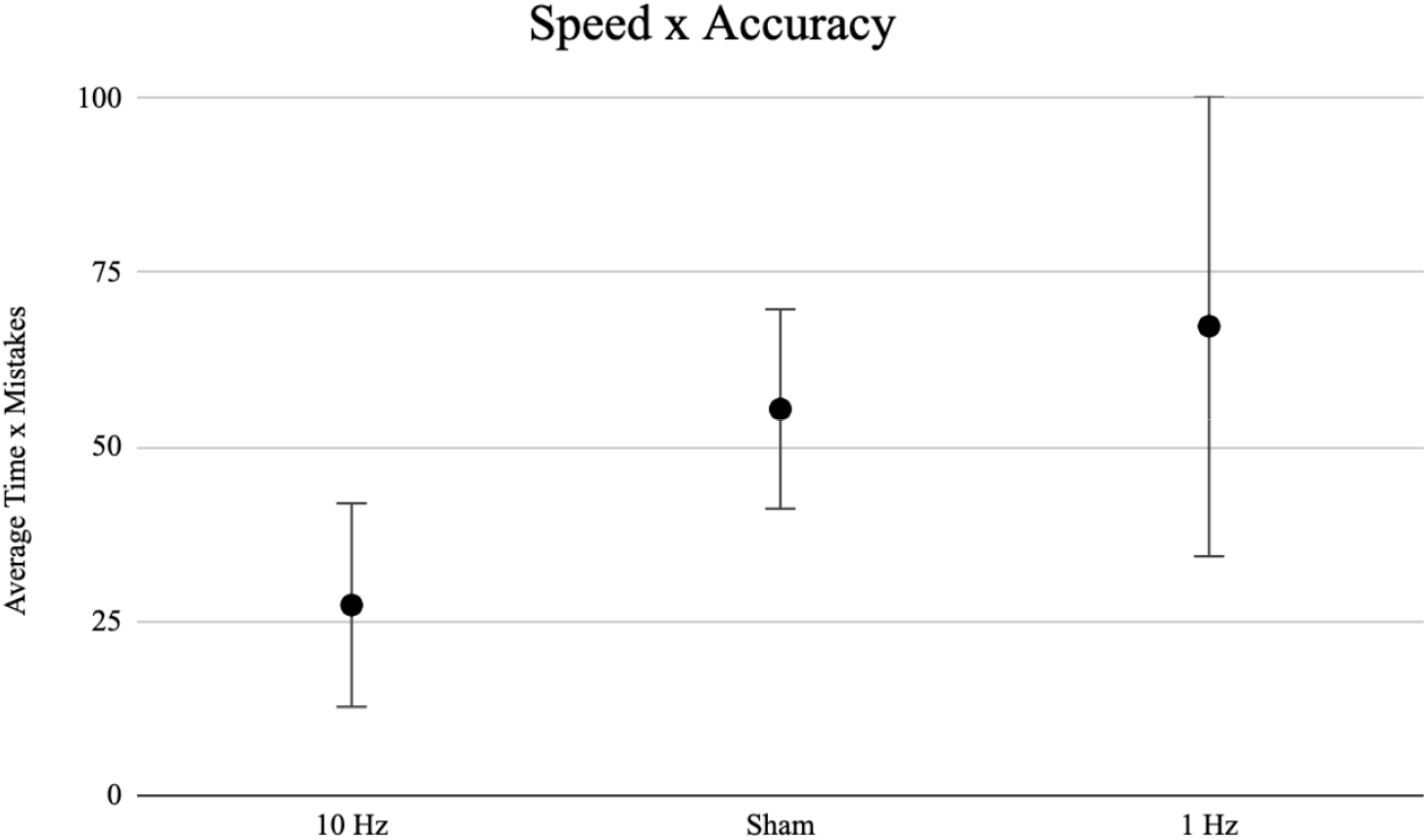
Combined measure of average speed multiplied by number of mistakes made for each TMS condition. All TMS conditions were found to be nonsignificant (p > 0.05). Standard errors of the means are plotted.

## 4. Discussion

This study demonstrates promising evidence that the rTPJ is responsible for at least part of a person’s ability to teach effectively. Despite the small sample size, a clear pattern was observed in which inhibiting the rTPJ caused a person to be a less efficient instructor, needing longer to complete each model, while exciting the rTPJ caused a person to be a more efficient instructor, needing less time to complete each model and improving the accuracy of the instruction. One of the main limitations of this study is its sample size. We were only able to recruit 5 participants, which strongly decreases the reliability of these results (Button et al., 2013). It is known that small sample sizes are a main contributor to type II errors in research analysis, which occurs when an effect truly exists, but an experiment is unable to show statistical significance and therefore incorrectly concludes that no effect exists (Mittendorf et al., 1995). A small sample size can also increase the chance of a type I error, in which an experiment incorrectly concludes that an effect exists (Leppink et al., 2016). Therefore, due to the sample size of the current study, it is impossible to truly draw any reliable conclusions.

Due to the known link between the rTPJ and a person’s ToM abilities (Boccadoro et al., 2019; Koster-Hale et al., 2017) in which TMS applied to the rTPJ directly influenced an individual’s ToM (Ahmad et al., 2021; LaVarco et al., 2022), we are assuming that the participants in this study experienced a similar change in ToM. Exciting the rTPJ should have caused an increase in ToM abilities (Sellaro et al., 2015), and while inhibiting the rTPJ should have had the opposite effect (Chou & Chen, 2021; Young, Camprodon, et al., 2010). It would then be consistent with the literature that the increased ToM abilities allowed participants to perform better on the Lego director task, and vice versa (Krych-Appelbaum et al., 2007). A flaw of the current study is that we did not directly measure the ToM levels of the participants during each TMS condition, so we instead assumed the change in ToM happened based on the literature. Future iterations of this study should incorporate a test such as the Minds in the Eye (MIE) test which would directly measure each participant’s ToM levels during each TMS condition. This would allow the conclusion to be made that the observed link between the rTPJ and a person’s teaching abilities is mediated by a person’s ToM abilities. With the study design as it is currently, we are unable to definitively tie these results to ToM.

In future iterations of this study, participants should also be given the Edinburgh Handedness Inventory to ensure that we are only using participants which are strongly right-handed. Evidence suggests that ToM is lateralized to the right hemisphere due to its association with the rTPJ (Harrison, 2016; Murray et al., 2021; Zevy & Cohen, 2016). Studies have found that an individual’s handedness is related to their neural mechanisms underlying lateralized processes in the brain (Propper et al., 2019; Sainburg, 2014). This study only used participants who were right-handed, but we did not assess whether they were strongly right-handed or if their preference was inconsistent.

In conclusion, excitation of the rTPJ improved an individual’s ability to teach efficiently and accurately. Given that TMS allows for causal establishment, it is apparent that teaching abilities are likely at least partially dependent on the rTPJ. Due to the link between the rTPJ and ToM abilities (Boccadoro et al., 2019; Dodell-Feder et al., 2011; Koster-Hale et al., 2017; Mossad et al., 2016), and the link between ToM abilities and teaching abilities (Achim et al., 2015; Champagne-Lavau et al., 2009; Krych-Appelbaum et al., 2007), it is reasonable to conclude that the relationship between the rTPJ and teaching abilities is mediated by ToM.

---

**Supplemental Table 1.** Analyses of Model Completion Times.

| Brain Stimulation Condition | Average Time to Complete a Model | SE | N |
| --- | --- | --- | --- |
| 10 Hz rTPJ | 32.93 s | 3.11 | 5 |
| 1 Hz rTPJ | 38.07 s | 2.76 | 5 |
| Sham | 34.03 s | 2.01 | 5 |

Hz, hertz; rTPJ, right temporoparietal junction; s, seconds; SE, standard error

**Supplemental Table 2.**
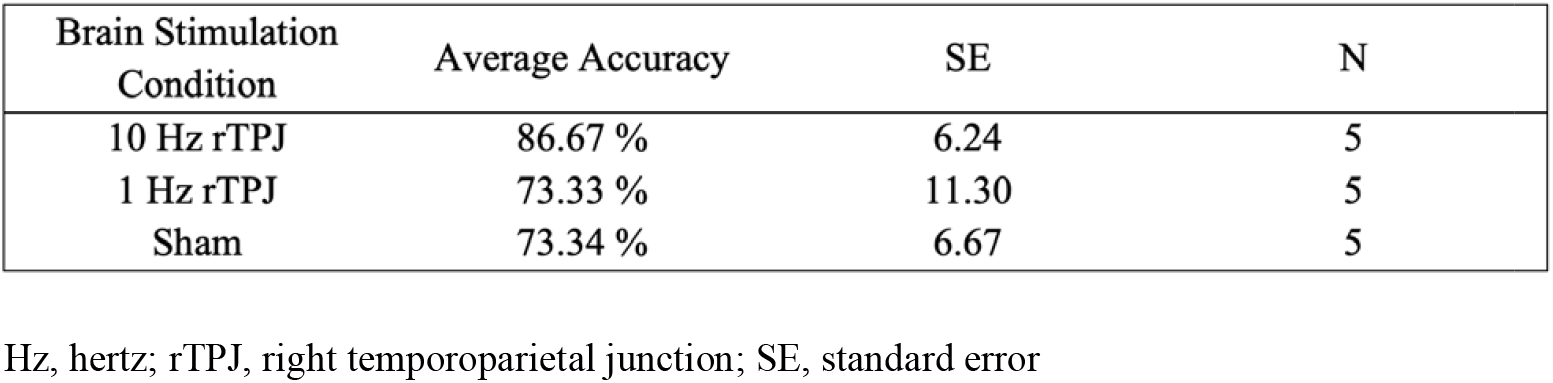
Analyses of Model Accuracy.

| Brain Stimulation Condition | Average Accuracy | SE | N |
| --- | --- | --- | --- |
| 10 Hz rTPJ | 86.67 % | 6.24 | 5 |
| 1 Hz rTPJ | 73.33 % | 11.30 | 5 |
| Sham | 73.34 % | 6.67 | 5 |

Hz, hertz; rTPJ, right temporoparietal junction; SE, standard error

**Supplemental Table 3.** Analyses of Combined Speed/Accuracy.

| Brain Stimulation Condition | Average Speed x Mistakes Value | SE | N |
| --- | --- | --- | --- |
| 10 Hz rTPJ | 27.34 | 14.59 | 5 |
| 1 Hz rTPJ | 67.33 | 33.05 | 5 |
| Sham | 55.46 | 14.31 | 5 |

Hz, hertz; rTPJ, right temporoparietal junction; SE, standard error

**Supplemental Table 4.** Time and Accuracy of Each Participant.

|  | Participant 1 |  |  | Participant 2 |  |  | Participant 3 |  |  | Participant 4 |  |  | Participant 5 |  |  |
| --- | --- | --- | --- | --- | --- | --- | --- | --- | --- | --- | --- | --- | --- | --- | --- |
|  | 10 Hz | Sham | 1 Hz | 10 Hz | Sham | 1 Hz | 10 Hz | Sham | 1 Hz | 10 Hz | Sham | 1 Hz | 10 Hz | Sham | 1 Hz |
| Model 1 | 27 s | 24 s | 37 s | 20 s | 22 | 40 | 27 s | 65 s | **52 s** | 32 s | 50 s | 42 s | 28 s | 22 s | 33 s |
| Model 2 | 27 s | 37 s | 28 s | **27 s** | 45 | 38 | 34 s | 25 s | 45 s | **32 s** | 33 s | 60 s | 34 s | 25 s | 33 s |
| Model 3 | 43 s | **52 s** | 31 s | 22 s | 50 | 39 | 48 s | 48 s | 70 s | 50 s | 76 s | 45 s | **55 s** | **35 s** | **39 s** |
| Model 4 | 25 s | 40 s | 47 s | 25 s | 23 | 24 | 40 s | 40 s | **70 s** | 71 s | **3 s** | 28 s | 24 s | **30 s** | 30 s |
| Model 5 | 26 s | 30 s | 28 s | 32 s | 19 | **40** | 52 s | **32 s** | **3 s** | 29 s | **28 s** | 21 s | 17 s | 28 s | **30 s** |
| Model 6 | 24 s | **17 s** | 26 s | 20 s | 31 | 27 | 39 s | **29 s** |  | **26 s** | 32 s | **44 s** | 32 s | 30 s | 44 s |
| Average (s) | 28.67 | 33.33 | 32.83 | 24.33 | 31.67 | 34.67 | 40.00 | 39.83 | 48.00 | 40.00 | 37.00 | 40.00 | 31.67 | 28.33 | 34.83 |
| Accuracy (%) | 100% | 66.67% | 100% | 83.33% | 100% | 83.33% | 100% | 66.67% | 33.33% | 66.67% | 66.67% | 83.33% | 83.33% | 66.67% | 66.67% |

Hz, hertz; s, seconds; bolded values indicate incorrect model

## References

Achim, A. M., Fossard, M., Couture, S., & Achim, A. (2015). Adjustment of speaker’s referential expressions to an addressee’s likely knowledge and link with theory of mind abilities. Frontiers in Psychology, 6, 823.

Ahmad, N., Zorns, S., Chavarria, K., Brenya, J., Janowska, A., & Keenan, J. P. (2021). Are We Right About the Right TPJ? A Review of Brain Stimulation and Social Cognition in the Right Temporal Parietal Junction. Symmetry, 13(11), 2219.

Apperly, I. A., Carroll, D. J., Samson, D., Humphreys, G. W., Qureshi, A., & Moffitt, G. (2010). Why are there limits on theory of mind use? Evidence from adults’ ability to follow instructions from an ignorant speaker. Q J Exp Psychol (Hove), 63(6), 1201–1217. 10.1080/17470210903281582

Bardi, L., Six, P., & Brass, M. (2017). Repetitive TMS of the temporo-parietal junction disrupts participant’s expectations in a spontaneous Theory of Mind task. Social Cognitive and Affective Neuroscience, 12(11), 1775–1782.

Baron-Cohen, S., Leslie, A. M., & Frith, U. (1985). Does the autistic child have a “theory of mind”? Cognition, 21(1), 37–46. 10.1016/0010-0277(85)90022-8

Baron-Cohen, S., Jolliffe, T., Mortimore, C., & Robertson, M. (1997). Another advanced test of theory of mind: evidence from very high functioning adults with autism or asperger syndrome. Journal of child psychology and psychiatry, and allied disciplines, 38(7), 813–822. 10.1111/j.1469-7610.1997.tb01599.x

Bitsch, F., Berger, P., Nagels, A., Falkenberg, I., & Straube, B. (2019). Impaired right temporoparietal junction–hippocampus connectivity in schizophrenia and its relevance for generating representations of other minds. Schizophrenia Bulletin, 45(4), 934–945.

Boccadoro, S., Cracco, E., Hudson, A. R., Bardi, L., Nijhof, A. D., Wiersema, J. R., Brass, M., & Mueller, S. C. (2019). Defining the neural correlates of spontaneous theory of mind (ToM): An fMRI multi-study investigation. Neuroimage, 203, 116193. 10.1016/j.neuroimage.2019.116193

Brunet-Gouet, E., & Decety, J. (2006). Social brain dysfunctions in schizophrenia: a review of neuroimaging studies. Psychiatry Research: Neuroimaging, 148(2-3), 75-92.

Buccino, G., Baumgaertner, A., Colle, L., Buechel, C., Rizzolatti, G., & Binkofski, F. (2007). The neural basis for understanding non-intended actions. Neuroimage, 36, T119–T127.

Button, K. S., Ioannidis, J. P., Mokrysz, C., Nosek, B. A., Flint, J., Robinson, E. S., & Munafò, M. R. (2013). Power failure: why small sample size undermines the reliability of neuroscience. Nature reviews neuroscience, 14(5), 365–376.

Carrington, S. J., & Bailey, A. J. (2009). Are there theory of mind regions in the brain? A review of the neuroimaging literature. Human brain mapping, 30(8), 2313–2335.

Chail, A., Saini, R. K., Bhat, P., Srivastava, K., & Chauhan, V. (2018). Transcranial magnetic stimulation: a review of its evolution and current applications. Industrial psychiatry journal, 27(2), 172.

Champagne-Lavau, M., Fossard, M., Martel, G., Chapdelaine, C., Blouin, G., Rodriguez, J.-P., & Stip, E. (2009). Do patients with schizophrenia attribute mental states in a referential communication task? Cognitive neuropsychiatry, 14(3), 217–239.

Chou, Y., & Chen, T.-Y. (2021). Disruption on right temporoparietal junction with transcranial magnetic stimulation affects moral judgment: No difference between first-and third-personal narration with TMS. Neuropsychologia, 157, 107858.

Clark, H. H., & Brennan, S. E. (1991). Grounding in communication.

Clark, H. H., Schreuder, R., & Buttrick, S. (1983). Common ground at the understanding of demonstrative reference. Journal of verbal learning and verbal behavior, 22(2), 245–258.

Correia, J., Formisano, E., Valente, G., Hausfeld, L., Jansma, B., & Bonte, M. (2014). Brain-based translation: fMRI decoding of spoken words in bilinguals reveals language-independent semantic representations in anterior temporal lobe. J Neurosci, 34(1), 332–338. 10.1523/JNEUROSCI.1302-13.2014

Csibra, G., & Gergely, G. (2006). Social learning and social cognition: The case for pedagogy. Processes of change in brain and cognitive development. Attention and performance XXI, 21, 249–274.

Dichter, G. S. (2022). Functional magnetic resonance imaging of autism spectrum disorders. Dialogues in clinical neuroscience.

Dodell-Feder, D., Koster-Hale, J., Bedny, M., & Saxe, R. (2011). fMRI item analysis in a theory of mind task. Neuroimage, 55(2), 705–712.

Esse Wilson, J., Trumbo, M. C., Wilson, J. K., & Tesche, C. D. (2018). Transcranial direct current stimulation (tDCS) over right temporoparietal junction (rTPJ) for social cognition and social skills in adults with autism spectrum disorder (ASD). Journal of Neural Transmission, 125, 1857–1866.

Fletcher-Watson, S., McConnell, F., Manola, E., & McConachie, H. (2014). Interventions based on the theory of mind cognitive model for autism spectrum disorder (ASD). Cochrane Database of Systematic Reviews, (3). 10.1002/14651858.cd008785.pub2

Filmer, H. L., Fox, A., & Dux, P. E. (2019). Causal evidence of right temporal parietal junction involvement in implicit theory of mind processing. Neuroimage, 196, 329–336.

Gallagher, H. L., & Frith, C. D. (2003). Functional imaging of ‘theory of mind’. Trends in cognitive sciences, 7(2), 77–83.

Gupta, R., Tranel, D., & Duff, M. C. (2012). Ventromedial prefrontal cortex damage does not impair the development and use of common ground in social interaction: implications for cognitive theory of mind. Neuropsychologia, 50(1), 145–152.

Gweon, H., Dodell□Feder, D., Bedny, M., & Saxe, R. (2012). Theory of mind performance in children correlates with functional specialization of a brain region for thinking about thoughts. Child development, 83(6), 1853–1868.

Harrison, A.-N. (2016). Exploring the Hemispheric Lateralization of Theory of Mind. Undergraduate Journal of Psychology at Berkley, 9.

Hillebrandt, H., Dumontheil, I., Blakemore, S. J., & Roiser, J. P. (2013). Dynamic causal modelling of effective connectivity during perspective taking in a communicative task. Neuroimage, 76, 116–124. 10.1016/j.neuroimage.2013.02.072

Hyde, D. C., Aparicio Betancourt, M., & Simon, C. E. (2015). Human temporal□parietal junction spontaneously tracks others’ beliefs: A functional near□infrared spectroscopy study. Human brain mapping, 36(12), 4831–4846.

Kana, R. K., Maximo, J. O., Williams, D. L., Keller, T. A., Schipul, S. E., Cherkassky, V. L., Minshew, N. J., & Just, M. A. (2015). Aberrant functioning of the theory-of-mind network in children and adolescents with autism. Molecular autism, 6, 1–12.

Keenan, J., Gallup Jr, G., & Falk, D. (2003). The face in the mirror: The search for the origin of consciousness. In: Harper Collins New York.

Keysar, B., Lin, S., & Barr, D. J. (2003). Limits on theory of mind use in adults. Cognition, 89(1), 25–41. 10.1016/s0010-0277(03)00064-7

Klein, H. S., Vanneste, S., & Pinkham, A. E. (2021). The limited effect of neural stimulation on visual attention and social cognition in individuals with schizophrenia. Neuropsychologia, 157, 107880.

Koster-Hale, J., Richardson, H., Velez, N., Asaba, M., Young, L., & Saxe, R. (2017). Mentalizing regions represent distributed, continuous, and abstract dimensions of others’ beliefs. Neuroimage, 161, 9–18.

Krall, S. C., Rottschy, C., Oberwelland, E., Bzdok, D., Fox, P. T., Eickhoff, S. B., Fink, G. R., & Konrad, K. (2015). The role of the right temporoparietal junction in attention and social interaction as revealed by ALE meta-analysis. Brain Structure and Function, 220, 587–604.

Krych-Appelbaum, M., Law, J. B., Jones, D., Barnacz, A., Johnson, A., & Keenan, J. P. (2007). “I think I know what you mean”: The role of theory of mind in collaborative communication. Interaction Studies, 8(2), 267–280.

LaVarco, A., Ahmad, N., Archer, Q., Pardillo, M., Nunez Castaneda, R., Minervini, A., & Keenan, J. P. (2022). Self-conscious emotions and the right fronto-temporal and right temporal parietal junction. Brain sciences, 12(2), 138.

Lee, S.-H., Kim, W., Chung, Y.-C., Jung, K.-H., Bahk, W.-M., Jun, T.-Y., Kim, K.-S., George, M. S., & Chae, J.-H. (2005). A double blind study showing that two weeks of daily repetitive TMS over the left or right temporoparietal cortex reduces symptoms in patients with schizophrenia who are having treatment-refractory auditory hallucinations. Neuroscience letters, 376(3), 177–181.

Leppink, J., Winston, K., & O’Sullivan, P. (2016). Statistical significance does not imply a real effect. Perspectives on medical education, 5, 122–124.

Mittendorf, R., Arun, V., & Sapugay, A. M. V. (1995). The problem of the type II statistical error. Obstetrics & Gynecology, 86(5), 857–859.

Mossad, S. I., AuCoin-Power, M., Urbain, C., Smith, M. L., Pang, E. W., & Taylor, M.J. (2016). Thinking about the thoughts of others; temporal and spatial neural activation during false belief reasoning. Neuroimage, 134, 320–327.

Mukerji, C. E., Lincoln, S. H., Dodell-Feder, D., Nelson, C. A., & Hooker, C. I. (2019). Neural correlates of theory-of-mind are associated with variation in children’s everyday social cognition. Social Cognitive and Affective Neuroscience, 14(6), 579–589.

Murray, E., Brenya, J., Chavarria, K., Kelly, K. J., Fierst, A., Ahmad, N., Anton, C., Shaffer, L., Kapila, K., & Driever, L. (2021). Corticospinal Excitability during a Perspective Taking Task as Measured by TMS-Induced Motor Evoked Potentials. Brain sciences, 11(4), 513.

Nijhof, A. D., Bardi, L., Brass, M., & Wiersema, J. R. (2018). Brain activity for spontaneous and explicit mentalizing in adults with autism spectrum disorder: An fMRI study. NeuroImage: Clinical, 18, 475–484.

Pobric, G., Lambon Ralph, M. A., & Zahn, R. (2016). Hemispheric specialization within the superior anterior temporal cortex for social and nonsocial concepts. Journal of Cognitive Neuroscience, 28(3), 351–360.

Powell, J. L., Grossi, D., Corcoran, R., Gobet, F., & Garcia-Finana, M. (2017). The neural correlates of theory of mind and their role during empathy and the game of chess: A functional magnetic resonance imaging study. Neuroscience, 355, 149–160.

Premack, D., & Woodruff, G. (1978). Does the chimpanzee have a theory of mind? Behavioral and brain sciences, 1(4), 515–526.

Propper, R. E., Wolfarth, A., Carlei, C., Brunye, T. T., & Christman, S. D. (2019). Superior categorical and coordinate spatial task performance in inconsistent-handers relative to consistent-right-handers. Laterality: Asymmetries of Body, Brain and Cognition, 24(3), 274–288.

Rodriguez, V. (2013). The Human Nervous System: A Framework for Teaching and the Teaching Brain. Mind, Brain, and Education, 7(1), 2–12. 10.1111/mbe.12000

Rubio-Fernandez, P. (2017). The director task: A test of Theory-of-Mind use or selective attention? Psychon Bull Rev, 24(4), 1121–1128. 10.3758/s13423-016-1190-7

Rubio-Fernandez, P. (2021). Pragmatic markers: the missing link between language and Theory of Mind. Synthese, 199(1-2), 1125–1158.

Sainburg, R. L. (2014). Convergent models of handedness and brain lateralization. Frontiers in Psychology, 5, 1092–1108.

Salehinejad, M. A., Paknia, N., Hosseinpour, A. H., Yavari, F., Vicario, C. M., Nitsche, M. A., & Nejati, V. (2021). Contribution of the right temporoparietal junction and ventromedial prefrontal cortex to theory of mind in autism: A randomized, sham□controlled tDCS study. Autism Research, 14(8), 1572–1584.

Saxe, R., & Kanwisher, N. (2013). People thinking about thinking people: the role of the temporo-parietal junction in “theory of mind”. In Social neuroscience (pp. 171–182). Psychology Press.

Saxe, R., & Powell, L. J. (2006). It’s the thought that counts: specific brain regions for one component of theory of mind. Psychological science, 17(8), 692–699.

Schober, M. F., Conrad, F. G., & Fricker, S. S. (2004). Misunderstanding standardized language in research interviews. Applied Cognitive Psychology, 18(2), 169–188.

Scholz, J., Triantafyllou, C., Whitfield-Gabrieli, S., Brown, E. N., & Saxe, R. (2009). Distinct regions of right temporo-parietal junction are selective for theory of mind and exogenous attention. PloS one, 4(3), e4869.

Schuwerk, T., Grosso, S. S., & Taylor, P. C. (2021). The influence of TMS of the rTPJ on attentional control and mentalizing. Neuropsychologia, 162, 108054.

Sellaro, R., Güro□lu, B., Nitsche, M. A., van den Wildenberg, W. P., Massaro, V., Durieux, J., Hommel, B., & Colzato, L. S. (2015). Increasing the role of belief information in moral judgments by stimulating the right temporoparietal junction. Neuropsychologia, 77, 400–408.

Suessbrick, A. L., Schober, M. F., & Conrad, F. G. (2000). Different respondents interpret ordinary questions quite differently. Proceedings of the American Statistical Association,

Van Overwalle, F., & Baetens, K. (2009). Understanding others’ actions and goals by mirror and mentalizing systems: a meta-analysis. Neuroimage, 48(3), 564–584.

Vélez, N., Chen, A. M., Burke, T., Cushman, F. A., & Gershman, S. J. (2023). Teachers recruit mentalizing regions to represent learners’ beliefs. Proceedings of the National Academy of Sciences, 120(22), e2215015120.

Völlm, B. A., Taylor, A. N., Richardson, P., Corcoran, R., Stirling, J., McKie, S., Deakin, J. F., & Elliott, R. (2006). Neuronal correlates of theory of mind and empathy: a functional magnetic resonance imaging study in a nonverbal task. Neuroimage, 29(1), 90–98.

Vucurovic, K., Caillies, S., & Kaladjian, A. (2020). Neural correlates of theory of mind and empathy in schizophrenia: An activation likelihood estimation meta-analysis. Journal of psychiatric research, 120, 163–174.

Wassermann, E. M. (1998). Risk and safety of repetitive transcranial magnetic stimulation: report and suggested guidelines from the International Workshop on the Safety of Repetitive Transcranial Magnetic Stimulation, June 5–7, 1996. Electroencephalography and Clinical Neurophysiology/Evoked Potentials Section, 108(1), 1–16.

Young, L., Camprodon, J. A., Hauser, M., Pascual-Leone, A., & Saxe, R. (2010). Disruption of the right temporoparietal junction with transcranial magnetic stimulation reduces the role of beliefs in moral judgments. Proceedings of the National Academy of Sciences, 107(15), 6753–6758.

Young, L., Dodell-Feder, D., & Saxe, R. (2010). What gets the attention of the temporo-parietal junction? An fMRI investigation of attention and theory of mind. Neuropsychologia, 48(9), 2658–2664.

Zevy, D., & Cohen, A. (2016). Hemispheric Lateralization of Theory of Mind. Western Undergraduate Psychology Journal, 4(1).

Zheng, L., Chen, C., Liu, W., Long, Y., Zhao, H., Bai, X., Zhang, Z., Han, Z., Liu, L., & Guo, T. (2018). Enhancement of teaching outcome through neural prediction of the students’ knowledge state. Human brain mapping, 39(7), 3046–3057.

